# Thermopriming-associated proteome and sugar content responses in *Pinus radiata* embryogenic tissue

**DOI:** 10.1101/2022.05.17.492240

**Authors:** Ander Castander-Olarieta, Cátia Pereira, Vera M. Mendes, Sandra Correia, Bruno Manadas, Jorge Canhoto, Itziar A. Montalbán, Paloma Moncaleán

**Affiliations:** Department of Forestry Science, NEIKER, Arkaute, Spain; Center for Functional Ecology, Department of Life Sciences, University of Coimbra, Coimbra, Portugal; CNC - Center for Neuroscience and Cell Biology, University of Coimbra, Coimbra, Portugal

**Author notes:** Correspondence: Paloma Moncaleán. These authors contributed equally as thesis co-directors (Itziar A. Montalbán, Paloma Moncaleán).

**Keywords:** carbohydrates, heat stress, metabolism, proteins, radiata pine, somatic embryogenesis

## Abstract

Improving the capacity of plants to face adverse environmental conditions requires a deep understanding of the molecular mechanisms governing stress response and adaptation. Proteomics, combined with metabolic analyses, offers a wide resource of information to be used in plant breeding programs. Previous studies have shown that somatic embryogenesis in *Pinus* spp. is a suitable tool not only to investigate stress response processes but also to modulate the behaviour of somatic plants. Based on this, the objective of this study was to analyse the protein and soluble sugar profiles of *Pinus radiata* embryonal masses after the application of high temperatures to unravel the mechanisms involved in thermopriming and memory acquisition at early stages of the somatic embryogenesis process. Results confirmed that heat provokes deep readjustments in the life-cycle of proteins, together with a significant reduction in the carbon-flux of central-metabolism pathways. Heat-priming also promotes the accumulation of proteins involved in oxidative stress defence, in the synthesis of specific amino acids such as isoleucine, influences cell division, the organization of the cytoskeleton and cell-walls, and modifies the levels of free soluble sugars like glucose or fructose. All this seems to be regulated by proteins linked with epigenetic, transcriptional and post-transcriptional mechanisms.

## 1 Introduction

Prompted by the current climate change situation, the production of elite plants with increased resilience to different adverse conditions has become a major concern during the last decades. In conifers, due to their large genome and their intrinsic heterozygosity, traditional breeding techniques have met limitations to fix desirable abiotic stress tolerance traits, hampering progress in forest-tree improvement (Williams and Svolainen 1996). Besides, these traits, as those from most agronomic species, are usually quantitative in nature, being regulated by loci widespread across the whole genome and exhibiting complex genetic control (Mosammaparast and Shi, 2010). In this sense, the emergence of epigenomics has opened up the avenue for a novel source of quick phenotypic variation to be exploited by breeders for the development of plants adapted to changing ecosystems (Samantara et al., 2021). In fact, epigenetics not only enables the modulation of qualitative traits but also the manipulation of intricate quantitative ones (Gahlaut et al., 2020).

Epigenetics, referred to as the mechanism by which heritable changes in gene expression occur without alterations in DNA sequence, involves the creation of a flexible memory that allows for a rapid and strong response to recurrent stress events. Epigenetic mechanisms control various developmental processes, plant growth and responses to environmental variations by regulating gene activity and fine-tuning gene expression (Amaral et al., 2020). These mechanisms, which may remain stable along plant life or across generations, provoke deep physiological and metabolic readjustments, including changes at the proteome level (Castander-Olarieta et al., 2021a). These kinds of alterations endow plants with the ability to showcase phenotypic plasticity and, hence, increase their adaptation capacity (Carbó et al., 2019).

Even when the role of stress memory is still supported by few researches in forestry species, probably due to the lack of fully sequenced genomes for most trees, there are multiple examples of experiments carried out during the last decade that reinforce this idea. This is the case, for example, of studies linking plasticity, drought stress response and epigenetics in poplar (Raj et al., 2011; Le Gac et al., 2018; Lafon-Placette et al., 2018). In this sense, although embryo development and seed maturation represent short periods of time when compared with the lifespan of a tree, they appear to be critical stages for the induction of phenotypic plasticity and the establishment of epigenetic memory, as reported in Norway spruce during both zygotic and somatic embryogenesis (SE) under different temperature conditions (Kvaalen and Johsen 2008; Yakovlev et al., 2010, 2011).

Multiple stress factors and environmental cues can induce persistent changes in epigenetic marks (Carbó et al., 2019). Particularly, heat stress is known to cause heritable phenotypic and epigenetic changes in several plant species such as *Arabidopsis thaliana* or wheat (Migicovsky et al. 2014; Wang et al., 2016), and priming and memory after exposure to heat stress have been proved in the past few years as an efficient experimental system to unravel the underlying molecular mechanisms (Oberkofler et al., 2021). Heat stress response is mainly mediated by heat shock proteins (HSPs) and heat shock factors, which act as molecular chaperones and transcription regulators, respectively. The former can activate a huge variety of enzymes, such as those involved in reactive oxygen species (ROS) scavenging and detoxification, or increase the levels of specific metabolites to be used for osmoregulation, signalling or membrane stabilisation, among others (Nishad and Nandi, 2021). Moreover, certain proteins have recently been associated with memory response because of their slow rate of turnover and persistence in the cells for long periods (Schneider et al., 2019), and specific soluble sugars such as glucose seem to have an indispensable role in the generation of thermomemory in *A. thaliana* (Sharma et al. 2019). In spite of this, a deeper understanding of the molecular machinery behind the process of thermopriming and thermomemory is required in woody plants.

In this regard, previous studies developed at out laboratory have shown that the application of high temperatures during initiation of both *Pinus radiata* and *P. halepensis* SE not only influence the success of the different stages of the process (Castander-Olarieta et al., 2019, 2021a; Pereira et al, 2020), but also the behaviour of the SE-derived plants years later under both standard and stress conditions (Castander-Olarieta et al., 2021b). These changes are associated with modifications in the profile of cytokinins (Castander-Olarieta et al., 2021c) and certain amino acids, in the pattern of epigenetic marks such as 5-hydroxymethylcytosine and 5-hydroxymethylcytosine, as well as in the expression of stress related genes (Castander-Olarieta et al., 2020; Pereira et al., 2021). On top of that, in our latest study it was found that thermopriming provoked a long-term reorganization of the proteome in radiata pine somatic embryos leading to alterations in sugar metabolism and translation mechanisms (Castander-Olarieta et al, 2021a). However, a complete image of how thermomemory is built and maintained has not yet been accomplished, and information regarding the involvement of proteins and metabolites at earlier stages of the process is needed.

To this aim, in this work we have analysed the proteome and soluble sugar content of proliferating radiata pine embryonal masses (EMs) after induction at high temperatures to assess whether the molecular and physiological differences observed at embryo and plant level are formed or could be derived from early alterations during EMs proliferation.

## 2 Materials and Methods

### 2.1 Plant material and temperature experiment

*Pinus radiata* D. Don green cones were collected in June 2018 from five genetically different mother trees in a seed orchard established by Neiker at Deba (Spain; latitude: 43°16’59”N, longitude: 2°17’59”W, altitude: 50 m). Cones were stored and processed as described in Montalbán et al. (2015) and seeds were sterilized following the same protocol. Megagametophytes containing immature zygotic embryos at polyembryony stage were excised out aseptically and placed horizontally onto initiation Embryo Development Medium (EDM, Walter et al., 2005), as described in Montalbán et al. (2012). For the temperature experiment, Petri plates (90 × 14 mm) containing EDM medium were prewarmed for 30 min and once the megagametophytes were placed onto the medium, they were cultured at 23 °C for 8 weeks (Cond1; control), 40 °C for 4 h (Cond2) and 60 °C for 5 min (Cond3). The experiment comprised 1200 megagametophytes (8 megagametophytes per Petri plate x 10 Petri plates x three treatments x five mother trees). After the treatments all plates were cultured at the standard conditions of 23 °C in darkness for 8 weeks. Then, actively proliferating EMs were detached from megagametophytes and subcultured to medium of the same composition. Fourteen days later, EMs were recorded as established cell lines (ECLs) and transferred to proliferation medium as described in Montalbán et al. (2012). After five subculturing periods on this medium (70 days), five EMs with different genetic background were selected per treatment and 500 mg of EMs were frozen in liquid nitrogen and stored at -80 °C until protein and sugar content analyses (Figure 1). Further steps of the embryogenic process as well as different measurements (initiation, proliferation and maturation rates) are described in Castander-Olarieta et al. (2021a).

**Fig. 1.**
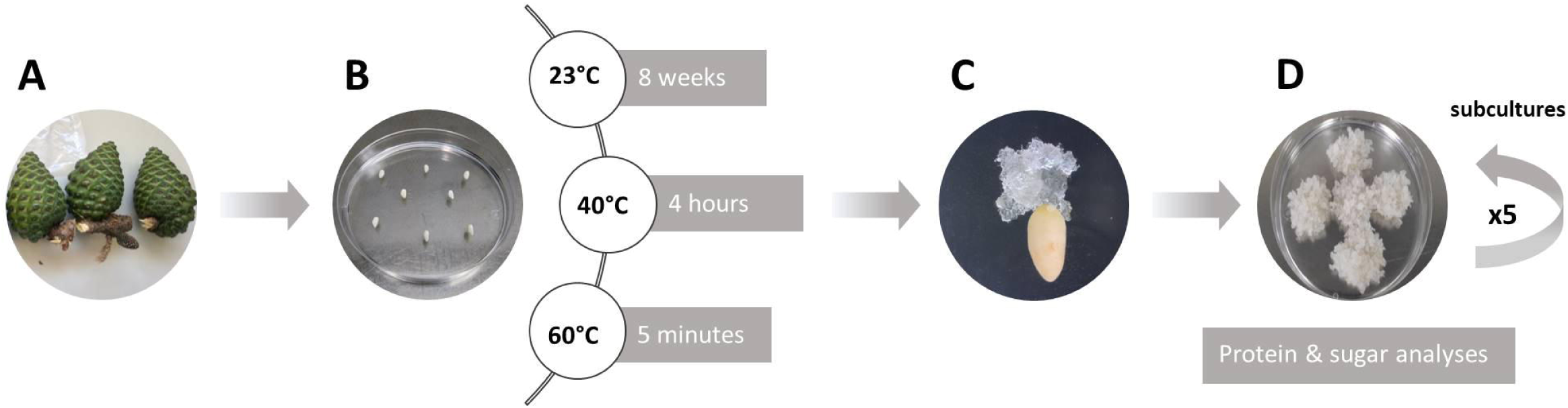
Schematic diagram of the experimental setup: (A) *P. radiata* green cones enclosing immature zygotic embryos at polyembryony stage from five genetically different mother trees; (B) Aseptically extracted megagametophytes cultured on EDM initiation medium under three temperature conditions (Cond1: 23 °C for 8 weeks; Cond2: 40 °C for 4 h; Cond3: 60 °C for 5 min); (C) Embryogenic tissue starting to grow out from megagametophytes after 8 weeks on culture medium; (D) Actively proliferating embryonal masses after five subculture periods on EDM proliferation medium ready to be frozen in liquid nitrogen for protein and soluble sugar analyses.

### 2.2. Protein and polar metabolite extraction

Protein and metabolite extractions from proliferating EMs were performed following the combined protocol described by Valledor et al. (2014) with slight modifications. Briefly, 2 mL of cold extraction buffer (methanol:chloroform:water 2.5:1:0.5, v:v:v) were added to 500 mg of liquid nitrogen grinded EMs. Five ECLs were used per temperature condition. Then, samples were centrifuged at 5000 g for 10 min at 4 °C.

The supernatant, containing metabolites, was transferred to a new microcentrifuge tube and 2 mL of phase separation mix (chroloform:water 1:1) were added. After centrifugation (5000 g, 10 min, room temperature), the upper aqueous phase was transferred again to a new tube for an extra fractionating step with 800 μL of phase separation mix. Finally, the upper layer containing polar metabolites was saved to new tubes and stored at -80 °C until HPLC analysis.

Pellets containing proteins and nuclei acids were washed with 2 mL of 0.75% (v/v) β-mercaptoethanol in 100% methanol, centrifuged (5000 g, 10 min, 4 °C) and the supernatant discarded. The washing step was repeated once. Pellets were then air dried and dissolved in 1 mL of pellet solubilisation buffer (Valledor et al., 2014) for incubation at 37 °C in a thermal shaker for 20 min. Then, samples were centrifuged (14000 g, 3 min) and the supernatants were transferred to silica columns to bind DNA (Zymo Research, Irvin, California, USA). After 1 min of incubation, columns were centrifuged (10000 g, 1 min) and the flowthrough was mixed with 600 μL of acetonitrile. The mix was transferred to a new silica column, incubated for 1 min, and centrifuged at 10000 g for 1 min. The flowthrough was transferred to a new tube and 1.1 mL of phenol and 1.2 mL of water were added. Samples were mixed, centrifuged (10000 g, 8 min) and the upper phenolic phase was transferred to a new tube that contained 1.2 mL PWB (Valledor et al., 2014). After vortexing and centrifugation (10000 g, 8 min, room temperature), upper phenolic phase was transferred to a new tube and proteins were precipitated overnight at -20 °C by the adding of 3 mL of 0.1 M ammonium acetate, 0.5% β-mercaptoethanol in methanol. Samples were then centrifuged at 10000 g for 20 min at 4 °C, the supernatant removed, and pellets were washed with cold acetone three times. Finally, proteins were precipitated by centrifugation (10000 g, 20 min) and pellets were air dried and re-suspended in 80 μL of solubilisation buffer [7M urea, 2M thiourea, 2% (w/v) CHAPS, 1% (w/v) DTT].

### 2.3 Protein sample preparation and LC-MS methodology

Aliquots of 70 µL were obtained from the protein extracts and precipitated using 280 µL of cold acetone for 30 min at -80 °C. Samples were then centrifuged at 20000 g, refrigerated at 4 °C for 20 min and the pellet re-suspended in 50 µL of 1× Laemmli Sample Buffer. The total protein concentration for each sample was measured using the Pierce 660 nm Protein Assay kit (Thermo Scientific™). At this point two different experiments were performed: an Ion-Library construction [data-dependent acquisition (DDA)] and the relative quantification of proteins (DIA; SWATH-MS). For DDA, all the samples from each condition were pooled, obtaining just one sample per treatment (Cond1, Cond2, and Cond3), and for DIA each sample was processed individually. Then, proteins from each sample were separated by SDS-PAGE for 17 min at 110 V (Short-GeLC Approach; Anjo et al., 2015) and stained with Coomassie Brilliant Blue G-250. In the case of DDA, each lane was divided into 5 gel pieces, whereas for DIA experiments lanes were divided into 3 gel pieces for further individual processing. Once destained, proteins were digested by overnight incubation with trypsin and peptides were extracted from the gel using increasing concentrations of acetonitrile (30, 50, and 98%) with 1% formic acid. Extraction solvent was evaporated using a vacuum-concentrator and peptides were re-suspended in 20 µL of 2% acetonitrile and 0.1% formic acid. Finally, samples were sonicated using a cup-horn (Ultrasonic processor, 750W) for 2 min, at 40% amplitude, and pulses of 1 sec ON/OFF.

For both DIA and DDA experiments, 10 µL of each sample were subjected to LC-MS/MS using a NanoLC™ 425 System (Eksigent) coupled to a Triple TOF™ 6600 mass spectrometer (Sciex) and the ionization source (ESI DuoSpray™ Source). The chromatographic separation was performed on a Triart C18 Capillary Column 1/32” (12 nm, S-3µm, 150 × 0.3 mm, YMC) and using a Triart C18 Capillary Guard Column (0.5 × 5 mm, 3 μm, 12nm, YMC). Further details of the LC methodology (mobile phases, flow rate etc) and the different mass spectrometer operation modes used for DDA and DIA experiments are resumed in Castander-Olarieta et al. (2021a). The only differences are the following: LC program [5 – 30% of B (0 - 50 min), 30 – 98% of B (50 – 52 min), 98% of B (52-54 min), 98 - 5% of B (54 – 56min), and 5% of B (56 – 65 min], voltage of the ionization source (5500 V), nebulizer gas 1 (GS1) and gas 2 (GS2) pressures (25 and 10 psi respectively), candidate ions minimum count threshold (100) and mass spectrometer software version (Analyst® TF 1.8.1, Sciex®).

### 2.4. Soluble sugar content analysis

Aliquots of 500 μL were obtained from the fractions containing polar metabolites in section 2.2 and the methanol of the extraction buffer was completely evaporated on a Speedvac. Metabolites were then re-suspended in 100 μL distilled water and soluble sugars and sugar alcohols (fructose, glucose, sucrose, mannitol, sorbitol) were quantified by HPLC (Agilent 1260 Infinity II) using an 8 μm Hi-Plex Ca column (7.7 mm x 300 mm, 8 µm) with a guard column and connected to a refractive index detector (RID). The mobile phase was pure water and the samples were injected in the column at a flow rate of 0.2 mL min^-1^ at 80 °C for 40 min. Sugar concentrations were determined from internal calibration curves constructed with the corresponding commercial standards and expressed as µmol g FW^−1^. The results were conveniently adjusted taking into account the concentration step (5 times) just after methanol evaporation and before HPLC analysis.

### 2.5 Data analysis

#### 2.5.1 Ion-Library construction (DDA information)

A specific ion-library of the precursor masses and fragment ions was created by combining all files from the DDA experiments in one protein identification search using the ProteinPilot™ software (v5.0, Sciex®). The paragon method parameters were the following: searched against the reviewed *Viridiplantae* database (Swissprot), cysteine alkylation by acrylamide, digestion by trypsin, and gel-based ID. Other reviewed databases and their combinations were also tested, such as the *Pinus taeda* database or the combination of *Solanaceae* + *Pinacea* + *Arabidopsis* + *Populus* databases and the *Solanaceae* + *Pinacea* + *Arabidopsis* + *Populus* + *Viridiplantae* databases. An independent False Discovery Rate (FDR) analysis, using the target-decoy approach provided by Protein Pilot™, was used to assess the quality of identifications.

#### 2.5.2 Relative quantification of proteins (SWATH-MS)

SWATH data processing was performed using SWATH™ processing plug-in for PeakView™ (v2.0.01, Sciex®). Protein relative quantification was performed in all samples using the information from the protein identification search. Quantification results were obtained for peptides with less than 1% of FDR and by the sum of up to 5 fragments/peptide. Each peptide was normalized for the total sum of areas for the respective sample. Protein relative quantities were obtained by the sum of the normalized values for up to 15 peptides/protein.

Finally, all the mass spectrometry proteomics data was deposited to the ProteomeXchange Consortium via the PRIDE (Perez-Riverol et al., 2019) partner repository with the data set identifier PXD030957.

### 2.6 Statistical analysis

Firstly, the relative quantification values obtained for all the samples from the SWATH-MS approach were subjected to a correlation analysis using the the Analysis ToolPak from Excel and the Pearson function to check whether those from the same condition followed the same behaviour.

To determine the effect of the temperature treatments on the protein profile of EMs, a cross-validation approach combining univariate and multivariate analyses as reported in previous studies (Castander-Olarieta et al., 2021a) was carried out. Using the MetaboAnalyst web-based platform (www.metaboanalyst.ca), we performed a partial least square-discriminant analysis (PLS-DA) to separate the three temperature conditions and identify the proteins contributing the most to that clustering based on variable importance in projection (VIP) values. Those proteins were then selected for a Gene Ontology (GO) enrichment analysis and were classified according to their biological function using the FunRich software and the Plants database from UniProt database (https://www.uniprot.org/help/plants). As univariate analysis, a Kruskall-Wallis test in combination with Dunn’s *post-hoc* analysis and Benjamini–Hochberg’s *p*-value adjustment was conducted. Proteins presenting *p* < 0.05 were compared with those showing VIP > 1 from the PLS-DA analysis and the ones in common were used to construct a hierarchical clustering heat map (Euclidean distance; Complete algorithm) using the MetaboAnalyst web-based platform.

For the levels of the different sugars analysed, we performed an analysis of variance (ANOVA) followed by Tukey’s test of Multiple Comparisons (α = 0.05) to assess in which comparisons among temperature treatments statistical differences were observed. When the ANOVA did not fulfil the normality hypothesis the Kruskal-Wallis test was performed.

## 3. Results

### 3.1 Relative quantification of proteins

DDA was performed to build the ion-library to be used in relative protein quantification. From the UniProtKB/Swissprot database, the number of reviewed protein sequences found for the species *Pinus taeda* was only 22; thus, the protein identification search was performed against the whole *Viridiplantae* database (40,256 protein sequences). The number of proteins identified using this database was 1133. The combination of other databases or the combination of the *Viridiplantae* database with the other databases did not increase the number of identified proteins. After considering 1% FDR and performing the quality filters previously mentioned in M&M section, protein relative quantification was obtained for 770 proteins. The correlation analysis between samples showed that samples were highly correlated between them, presenting values higher than 0.92 for comparisons among samples from different temperature conditions, and even higher (> 0.96) for samples belonging to the same temperature condition (Supplementary Figure 1).

In order to reduce the dimensionality of the dataset generated during protein quantification and identify those proteins contributing to the discrimination between temperature treatments, a supervised multivariate analysis (PLS-DA) was used. The score plot for this analysis is shown in Figure 2A, with component 1 describing 19.1% of the variance and component 2 describing 18.7%. The PLS-DA performance was evaluated by means of cross-validation for the maximum number of four components, using the 10-fold method. According to the guidepost (Q^2^), an evaluation method of the predictivity ability of the model, the better prediction performance was provided by the model with two components (Figure 2C). The score plots of these two components clearly grouped and differentiated among the three temperature conditions, without overlap, as observed in Figure 2B by applying the third component. Figure 2A also indicated a major effect of the treatments in component 1, while the second component explained mainly differences among samples or ECLs.

**Fig. 2.**
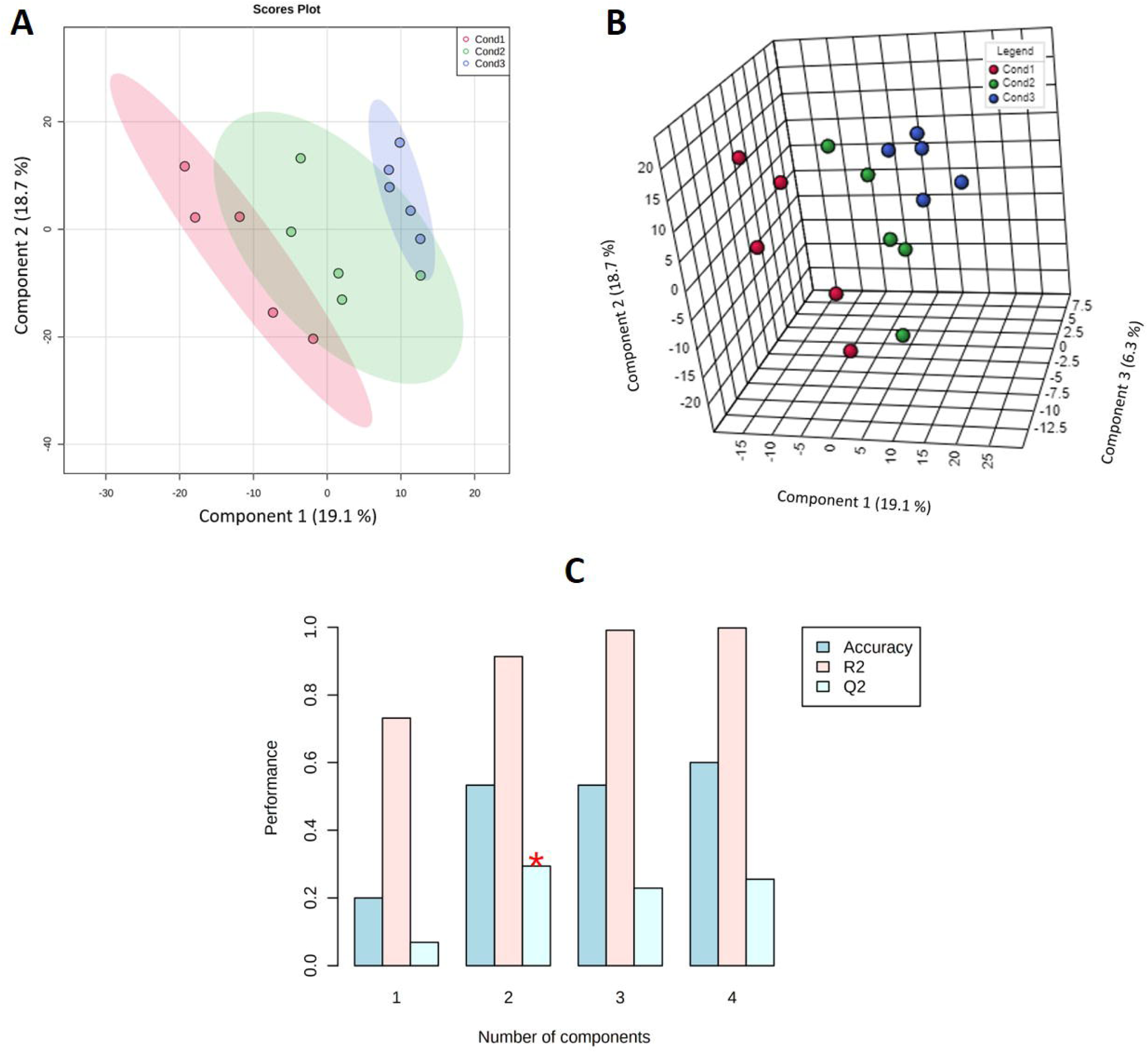
(A) Representation of the scores of the two first components for the PLS-DA analysis showing the ovals at 95% confidence interval using the 770 quantified proteins extracted from embryonal masses of *P. radiata* induced under high temperature conditions (Cond1 = 23 °C, control; Cond2 = 40 °C, 4 h; Cond3 = 60 °C, 5 min). (B) Representation of the scores of the three first components for the same PLS-DA analysis. (C) Results for the cross-validation of the PLS-DA analysis for the maximum number of four components by applying the 10-fold method. The red star indicates the best classifier according to the guidepost (Q^2^).

The VIP values obtained were chosen as criteria for selecting the most important variables on the PLS-DA model. Thus, only those proteins with a VIP value exceeding one were considered differentiating and the ones contributing to the separation between the three temperature conditions. This ranking included 278 proteins (36% of the total number of proteins used in the quantitative analysis (Supplementary Table 1), that were selected to perform a gene ontology enrichment analysis based on their biological function. This analysis revealed that the biological functions “response to cadmium ion”, “glycolytic process”, “proteasome-mediated ubiquitin-dependent protein catabolic process”, “tricarboxylic acid cycle”, “proteasome assembly” and “protein folding” were significantly over-represented (p < 0.05), and the percentage of proteins with those functions ranged from 1.6% to 8.8% in our study (Figure 3).

**Fig. 3.**
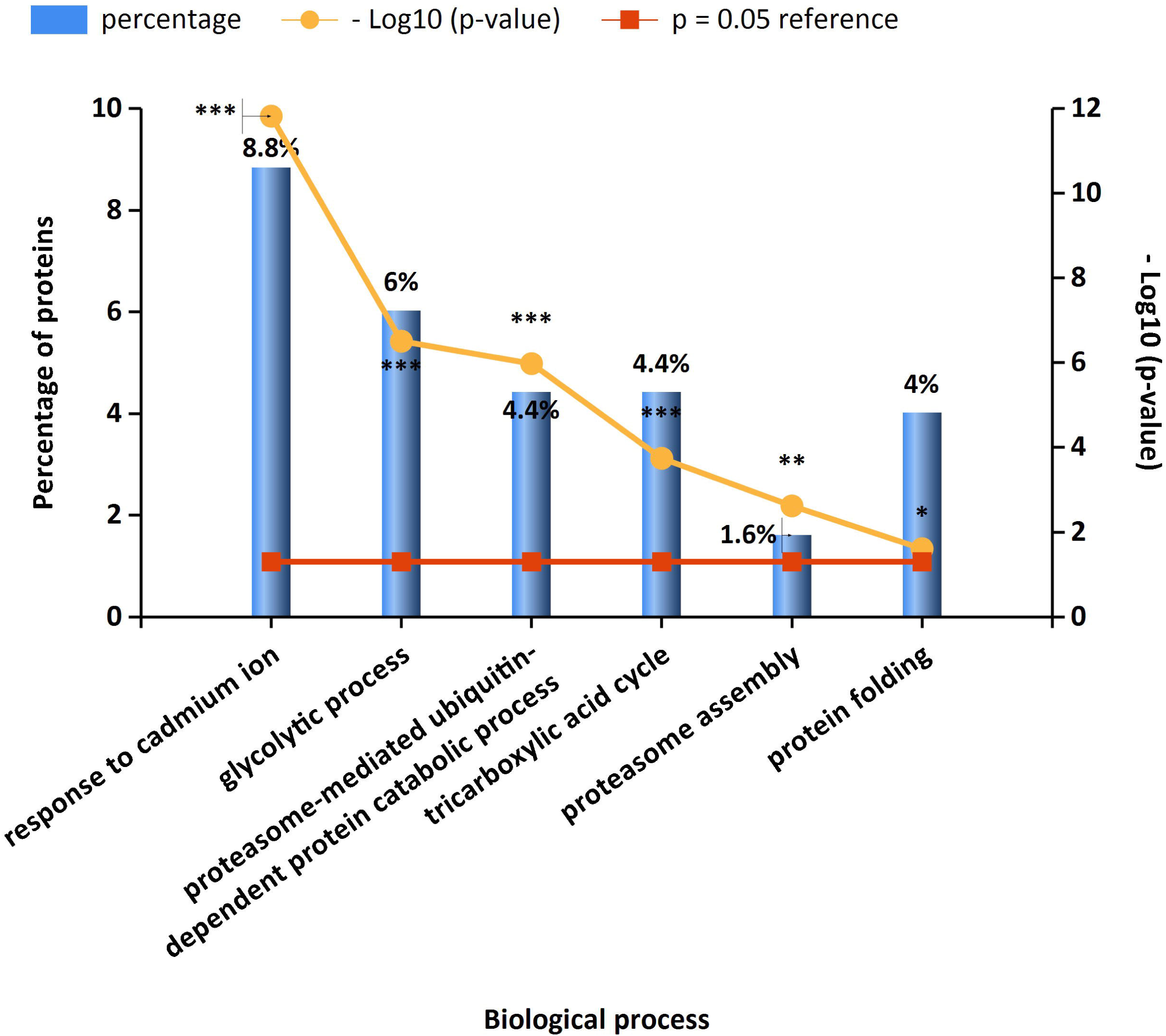
Gene ontology enrichment analysis of the 278 proteins selected from the PLS-DA analysis presenting VIP values greater than 1. The biological function clustering was performed using the FunRich software and the Plants database from UniProtdatabase. For each biological function the percentage of proteins belonging to that group as well as their significance level (*p* < 0.05) is indicated.

To double-check our results, a univariate statistical analysis was performed, showing that 68 proteins presented differential accumulation (*p* < 0.05) among the three conditions assayed. These proteins were then compared with the ones selected from the multivariate analysis, and the overlapping ones (53), that is, the ones resulting from the application of the cut-off of PLS-DA VIP score > 1 and *p* < 0.05, were considered the top significant ones or those most directly involved in heat-stress response and thermopriming (Figure 4). Proteins selected by this approach and their accession numbers, fold-changes between conditions, and VIP and *p* values are shown in Tables 1 and 2.

**Table 1.** Top significant proteins selected from the combination of the Kruskal-Wallis test and the PLS-DA analysis (p < 0.05 and VIP > 1) in embryonal masses of *P. radiata* induced under high temperature conditions (Cond1 = 23 °C, control; Cond2 = 40 °C, 4 h; Cond3 = 60 °C, 5 min).

| Description | Accession | Fold Change |  |  | Kruskal-Wallis | PLS-DA |
| --- | --- | --- | --- | --- | --- | --- |
|  |  | Cond2/Cond1 | Cond3/Cond1 | Cond3/Cond2 | p-values | VIP |
| 40S ribosomal protein S4-1 | Q93VH9 | 0.4 | 0.1 | 0.2 | 0.002 | 2.64 |
| Peptide methionine sulfoxide reductase | Q9SEC2 | 0.6 | 0.5 | 0.8 | 0.005 | 1.82 |
| Uncharacterized oxidoreductase At1g06690 | Q94A68 | 0.6 | 0.8 | 1.3 | 0.006 | 1.27 |
| Tubulin alpha-1 chain | Q43473 | 0.6 | 0.5 | 0.8 | 0.007 | 2.54 |
| Cellulose synthase-like protein D3 | Q9M9M4 | 0.7 | 0.6 | 0.9 | 0.008 | 2.25 |
| Succinate--CoA ligase [ADP-forming] subunit beta | Q84LB6 | 0.5 | 0.5 | 0.9 | 0.009 | 2.29 |
| Isocitrate dehydrogenase [NAD] regulatory subunit 3 | O81796 | 0.7 | 0.7 | 1.0 | 0.009 | 2.11 |
| Ketol-acid reductoisomerase | Q65XK0 | 1.1 | 1.7 | 1.5 | 0.010 | 2.28 |
| 3-oxoacyl-[acyl-carrier-protein] synthase I | P52410 | 1.4 | 1.7 | 1.2 | 0.011 | 2.39 |
| ATP-dependent zinc metalloprotease FTSH 2 | Q655S1 | 0.6 | 0.5 | 0.8 | 0.011 | 2.17 |
| S-adenosylmethionine synthase | Q944U4 | 1.1 | 1.8 | 1.7 | 0.012 | 2.32 |
| 6-phosphogluconate dehydrogenase, decarboxylating 2 | Q9FWA3 | 0.7 | 0.7 | 1.1 | 0.014 | 1.76 |
| 26S proteasome non-ATPase regulatory subunit 8 homolog A | Q9SGW3 | 0.8 | 1.3 | 1.7 | 0.015 | 1.36 |
| Transketolase-1 | Q8RWV0 | 0.7 | 1.5 | 2.2 | 0.016 | 1.24 |
| Glutamate--cysteine ligase | Q9ZNX6 | 0.7 | 0.7 | 1.1 | 0.018 | 1.71 |
| Rop guanine nucleotide exchange factor 7 | Q9LZN0 | 0.2 | 0.2 | 1.3 | 0.018 | 1.03 |
| Thioredoxin H-type | O65049 | 1.0 | 1.7 | 1.8 | 0.018 | 1.91 |
| Mitochondrial import inner membrane translocase subunit TIM9 | Q9XGX9 | 0.9 | 1.7 | 2.0 | 0.019 | 1.66 |
| Probable aldo-keto reductase 4 | Q93ZN2 | 0.9 | 0.5 | 0.6 | 0.021 | 2.05 |
| Zinc finger CCCH domain-containing protein 31 | Q7XPK1 | 0.7 | 0.7 | 1.0 | 0.021 | 1.84 |
| Plant UBX domain-containing protein 8 | F4JPR7 | 0.6 | 1.6 | 2.5 | 0.022 | 1.38 |
| Phosphoenolpyruvate carboxylase 1 | P29195 | 0.3 | 0.5 | 1.7 | 0.022 | 1.43 |
| Glutathione S-transferase DHAR3 | Q8LE52 | 1.2 | 1.5 | 1.2 | 0.024 | 2.13 |
| Stromal 70 kDa heat shock-related protein | Q02028 | 0.5 | 0.5 | 1.0 | 0.026 | 1.87 |
| UDP-arabinopyranose mutase 1 | Q9SRT9 | 1.3 | 1.4 | 1.1 | 0.032 | 1.69 |
| 40S ribosomal protein S20 | P35686 | 1.5 | 2.7 | 1.8 | 0.032 | 2.28 |
| RuBisCO large subunit-binding protein subunit beta | P08927 | 0.7 | 2.5 | 3.6 | 0.032 | 1.62 |

**Table 2.** Continuation of Table 1.

| Description | Accession | Fold Change |  |  | Kruskal-Wallis | PLS-DA |
| --- | --- | --- | --- | --- | --- | --- |
|  |  | Cond2/Cond1 | Cond3/Cond1 | Cond3/Cond2 | p-values | VIP |
| Ubiquinol oxidase 1a | O82807 | 0.5 | 0.5 | 1.1 | 0.032 | 1.76 |
| 60S ribosomal protein L23 | P49690 | 1.3 | 2.0 | 1.5 | 0.035 | 2.01 |
| NADH dehydrogenase [ubiquinone] flavoprotein 1 | Q9FNN5 | 1.0 | 0.7 | 0.7 | 0.036 | 1.49 |
| Mitochondrial dicarboxylate/tricarboxylate transporter DTC | Q9C5M0 | 0.6 | 0.5 | 0.7 | 0.037 | 1.94 |
| Cytochrome b-c1 complex subunit 7-1 | Q9SUU5 | 0.9 | 1.8 | 2.1 | 0.037 | 1.59 |
| Pathogenesis-related thaumatin-like protein 3.8 | A4PBQ1 | 0.5 | 1.9 | 4.1 | 0.037 | 1.08 |
| NADH dehydrogenase [ubiquinone] iron-sulfur protein 3 | Q37787 | 0.8 | 0.7 | 0.9 | 0.039 | 2.06 |
| Multiple organellar RNA editing factor 8 | Q9LKA5 | 0.9 | 1.2 | 1.3 | 0.039 | 1.35 |
| 26S proteasome regulatory subunit 4 homolog A | Q9SZD4 | 0.5 | 0.6 | 1.3 | 0.039 | 1.41 |
| Malate dehydrogenase 1 | P93819 | 0.6 | 0.6 | 1.0 | 0.041 | 1.68 |
| Glutamate dehydrogenase | P93541 | 0.6 | 0.6 | 0.9 | 0.042 | 1.99 |
| Formate dehydrogenase | Q9S7E4 | 1.3 | 2.0 | 1.6 | 0.042 | 1.94 |
| Ribonucleoside-diphosphate reductase large subunit | Q9SJ20 | 0.4 | 0.7 | 1.7 | 0.042 | 1.01 |
| Enolase | Q43130 | 0.6 | 0.4 | 0.7 | 0.043 | 1.87 |
| T-complex protein 1 subunit gamma | Q84WV1 | 0.5 | 0.7 | 1.5 | 0.044 | 1.04 |
| Casein kinase II subunit alpha-1 | Q08467 | 0.7 | 0.8 | 1.1 | 0.044 | 1.52 |
| Triosephosphate isomerase | P48496 | 0.7 | 0.6 | 0.9 | 0.046 | 1.96 |
| Splicing factor 3B subunit 6-like protein | Q9FMP4 | 0.4 | 0.4 | 1.0 | 0.046 | 1.79 |
| OVARIAN TUMOR DOMAIN-containing<br>deubiquitinating enzyme 2 | Q9LPT6 | 1.1 | 0.8 | 0.7 | 0.046 | 1.22 |
| Peptidyl-prolyl cis-trans isomerase CYP19-2 | Q9SKQ0 | 1.0 | 1.3 | 1.3 | 0.048 | 1.50 |
| 40S ribosomal protein S19-3 | Q9FNP8 | 1.2 | 2.5 | 2.1 | 0.048 | 1.99 |
| V-type proton ATPase subunit F | Q9ZQX4 | 0.8 | 0.8 | 1.1 | 0.048 | 1.16 |
| Importin subunit alpha-1a | Q71VM4 | 1.1 | 1.4 | 1.2 | 0.049 | 1.76 |
| 26S proteasome non-ATPase regulatory subunit<br>11 homolog | Q9LP45 | 0.5 | 0.5 | 1.1 | 0.049 | 1.70 |
| NAD-dependent malic enzyme 2 | Q8L7K9 | 0.6 | 0.5 | 0.9 | 0.050 | 1.87 |
| Glucose-1-phosphate adenylyltransferase small<br>subunit | Q9M462 | 0.5 | 0.5 | 1.0 | 0.050 | 1.44 |

**Fig. 4.**
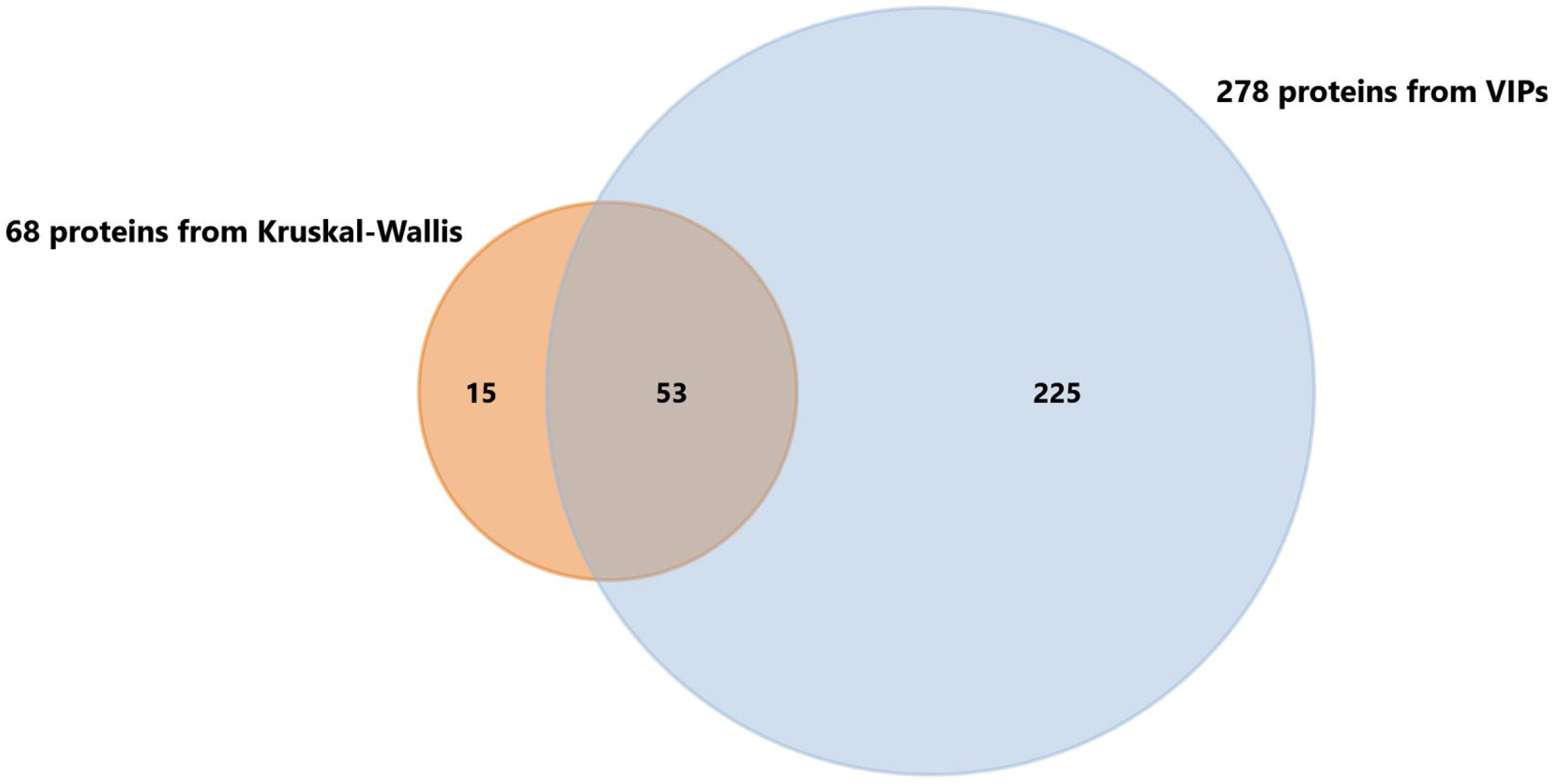
Venn-diagram showing that 53 proteins were common between the 68 with *p* < 0.05 from the Kruskal-Wallis test and the 278 with VIP > 1 from the PLS-DA analysis.

These proteins were finally used to build a heatmap plot (Figure 5). This heatmap plot shows to main blocks of proteins: the first block (above the yellow line) comprises a group of proteins whose concentration increases in EMs initiated under 60 °C for 5 min (Cond3), whereas their levels at 40 °C for 4 h (Cond2) remain similar to the ones of the control treatment of 23 °C (Cond1). However, the proteins in the second block presented a similar behaviour for both high temperature treatments (Cond2 and Cond3), showing lower levels than in the control treatment.

**Fig. 5.**
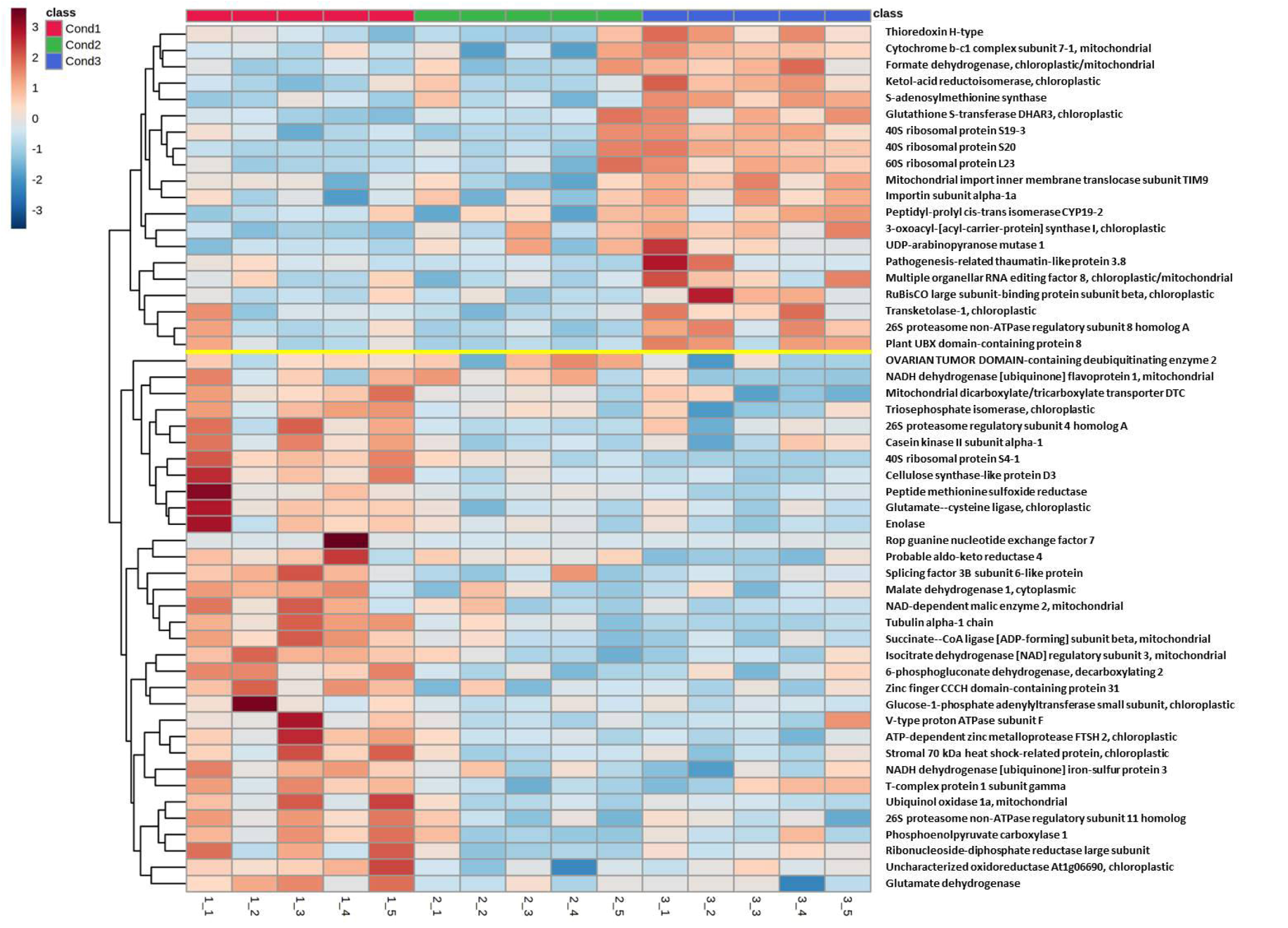
Hierarchical clustering heatmap using the 53 proteins having *p* < 0.05 and VIP score > 1 in embryonal masses of radiata pine induced under high temperature conditions (Cond1 = 23 °C, control; Cond2 = 40 °C, 4 h; Cond3 = 60 °C, 5 min). Hierarchical clustering was performed at the protein (rows) using Euclidean distance and Complete for the clustering algorithm. The yellow line separates the two blocks of proteins showing contrasting behaviour among treatments.

The first block (Figure 5) identifies a group of proteins involved in the life-cycle of proteins, such as those directly related with translation: some structural constituents of ribosomes like 40S ribosomal proteins S19-3 and S20, and 60S ribosomal protein L23. Besides, these proteins presented high fold changes between the control treatment (Cond1) and the highest temperature (Cond3). Other proteins in this group are involved in protein degradation (26S proteasome non-ATPase regulatory subunit 8 homolog A), in protein transport and folding (mitochondrial import inner membrane translocase subunit TIM9) and in accelerating protein folding (peptidyl-prolyl cis-trans isomerase CYP19-2). Following the same behaviour (increasing under Cond3) it was also detected proteins implicated in oxidative response and redox homeostasis (thioredoxin H-type; glutathione S-transferase DHAR3), in the synthesis of cell wall non-cellulosic polysaccharides, isoleucine and fatty acids (UDP-arabinopyranose mutase 1; ketol-acid reductoisomerase; 3-oxoacyl-[acyl-carrier-protein] synthase I; respectively), in methylation and RNA editing (S-adenosylmethionine synthase; multiple organellar RNA editing factor 8), one enzyme from the pentose phosphate pathway (transketolase-1) and one component of the electron transport chain (cytochrome b-c1 complex subunit 7-1). Finally, we have to remark the presence of other two proteins presenting high fold changes between Cond1 and Cond3 (Tables 1 and 2): a protein involved in abiotic stress response (formate dehydrogenase) and in the assembly of the enzyme RuBisCO (RuBisCO large subunit-binding protein subunit beta).

In the second block (Figure 5) it was also identified a large group of proteins related with protein synthesis and degradation (40S ribosomal protein S4-1; ATP-dependent zinc metalloprotease FTSH 2; 26S proteasome regulatory subunit 4 homolog A; casein kinase II subunit alpha-1; 26S proteasome non-ATPase regulatory subunit 11 homolog). Some of them also presented high fold changes between treatments, such as 40S ribosomal protein S4-1, a structural component of ribosomes (Tables 1 and 2). One of the biggest groups in this block, showing lower levels under both Cond2 and Cond3, is the one formed by central-metabolism enzymes, e.g., tricarboxylic acid cycle or glycolysis (succinate--CoA ligase [ADP-forming] subunit beta; isocitrate dehydrogenase [NAD] regulatory subunit 3; 6-phosphogluconate dehydrogenase, decarboxylating 2; phosphoenolpyruvate carboxylase 1; malate dehydrogenase 1; glutamate dehydrogenase; enolase; triosephosphate isomerase; NAD-dependent malic enzyme 2). It is worth highlighting the high fold changes of enolase and phosphoenolpyruvate carboxylase 1 (Tables 1 and 2). Other group of proteins in this block included oxidative stress-response enzymes (peptide methionine sulfoxide reductase; glutamate--cysteine ligase; probable aldo-keto reductase 4; ubiquinol oxidase 1a; V-type proton ATPase subunit F), proteins involved in cell division, cytoskeleton and cell-wall organization (tubulin alpha-1 chain; Rop guanine nucleotide exchange factor 7; ribonucleoside-diphosphate reductase large subunit; T-complex protein 1 subunit gamma), with high fold changes observed for Rop guanine nucleotide exchange factor 7 and ribonucleoside-diphosphate reductase large subunit (Tables 1 and 2), enzymes taking part in the synthesis of cell-wall and reserve polysaccharides (cellulose synthase-like protein D3; glucose-1-phosphate adenylyltransferase small subunit), some proteins from the electron transport chain (NADH dehydrogenase [ubiquinone] flavoprotein 1; NADH dehydrogenase [ubiquinone] iron-sulfur protein 3), proteins involved in gene expression regulation and RNA splicing, presenting the formers high fold changes (Tables 1 and 2) (zinc finger CCCH domain-containing protein 31; splicing factor 3B subunit 6-like protein; respectively), a heat shock protein (stromal 70 kDa heat shock-related protein) and one protein related with fatty acid transport and elongation (mitochondrial dicarboxylate/tricarboxylate transporter DTC).

### 3.2 Soluble sugar content quantification

The sugar content analysis using the HPLC methodology enabled the quantification of four (glucose, fructose, sucrose and sorbitol) of the five sugars and sugar alcohols tested. The two most abundant sugars were glucose and fructose respectively, presenting concentrations of one order of magnitude higher than the third most abundant sugar, the disaccharide sucrose. In comparison with these sugars, the sugar alcohol sorbitol was found at very low concentrations, presenting values around 0.04 µmol g^-1^ FW. Taking into account the effect of the treatments, significant differences were observed for fructose and glucose (*p* < 0.05). The levels of these two monosaccharides were considerably increased at 40 °C for 4 h treatment (Cond2) (Figure 6). In the case of sucrose, despite not significant, the opposite behaviour was detected, being the treatment at 40 °C for 4 h the one showing the lowest concentrations.

**Fig. 6.**
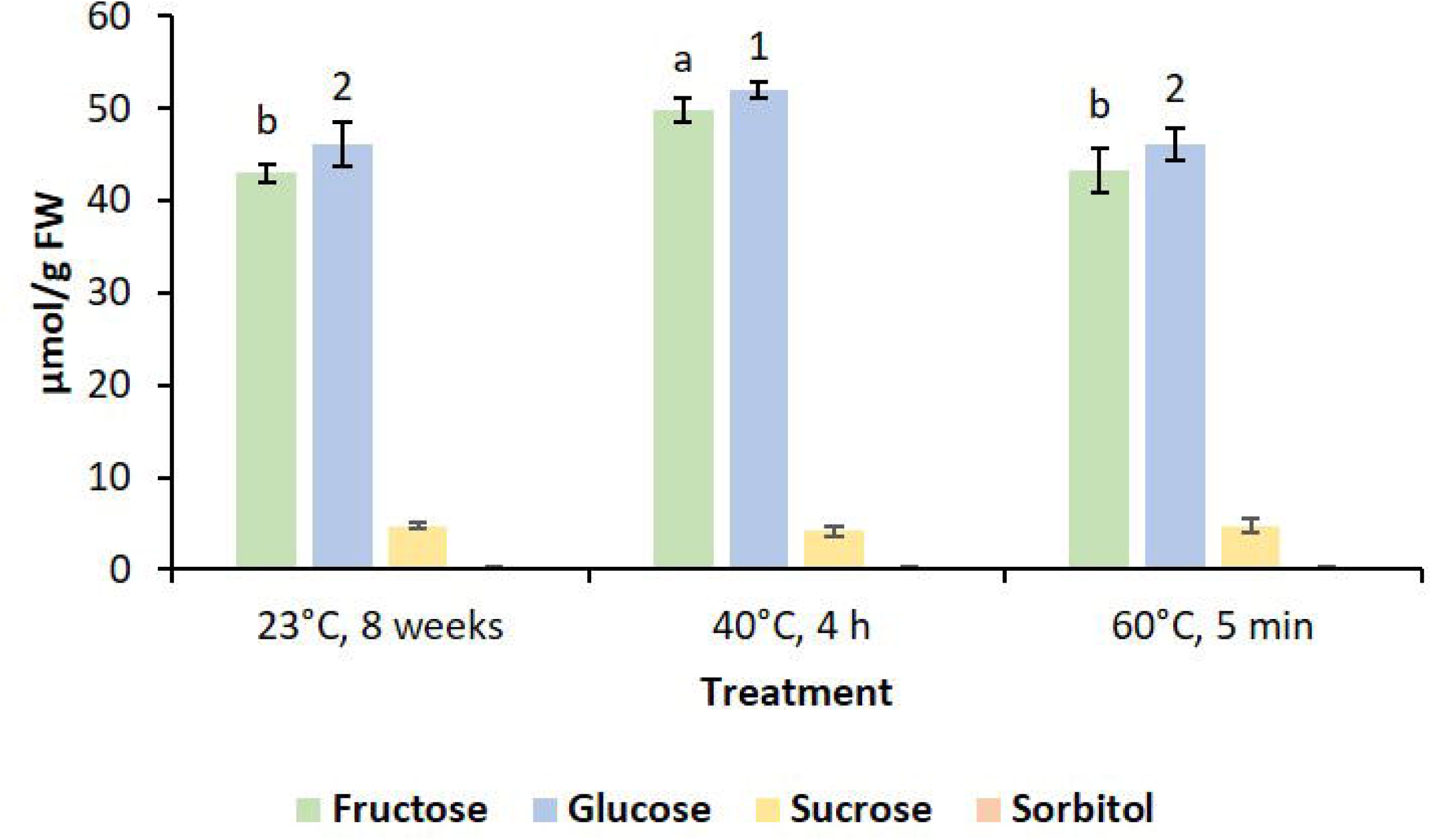
Effect of temperature treatments during induction of somatic embryogenesis (23 °C, 8 weeks; 40 °C, 4 h; 60 °C, 5 min) on the levels of fructose, glucose, sucrose and sorbitol in proliferating embryonal masses of *P. radiata*. Data are presented as mean values ± SE. Significant differences among treatments at *p* < 0.05 were analysed by Tukey’s test of Multiple Comparisons and were only found for fructose (indicated by different letters) and glucose (indicated by different numbers).

## 4. Discussion

Understanding the mechanisms governing plant stress response and memory acquisition is the first step towards the possibility of modulating the behaviour of plants and improving breeding efficiency. In this regard, in this work proteome analysis was used, together with soluble sugar and sugar alcohol determination, to discover new horizons involved in heat stress response and memory acquisition/maintenance at early stages of the SE process in an economically important tree species such as radiata pine. To this aim, a short-gel-liquid chromatography coupled to tandem mass spectrometry (Short-GeLCMS/MS) approach was employed, using DDA and DIA acquisition methods, in thermoprimed vs control EMs. This methodology enabled the identification of 1133 proteins; average values if compared with the ones from previous studies in radiata pine plants (Escandón et al., 2017; Pascual et al., 2017; 901 and 1228 proteins, respectively) or embryogenic cultures from other conifer species (dos Santos et al., 2016; Douétts-Peres et al., 2019; 2398 and 1518 proteins respectively). These values are also in accordance with the ones obtained in our previous work in radiata pine somatic embryos (Castander-Olarieta et al., 2021a; 1020 proteins). We have to highlight that this is, as far as we know, the first report of proteomics in EMs of this tree species.

The combination of multivariate statistical analysis and the GO enrichment analysis revealed that a big group of proteins responded distinctly to the three temperature conditions. Among them, some biological functions were significantly over-represented, such as those related to central metabolism and sugar catabolism (glycolysis and tricarboxylic acid cycle), protein folding, assembly and catabolism, and response to cadmium ion. Similar results were obtained in our previous work carried out with somatic embryos in which we detected a high number of proteins involved in the life-cycle of proteins and in sugar metabolism (Castander-Olarieta et al., 2021a). The presence of proteins related to response to cadmium stress is somehow surprising, as no cadmium was added to our culture medium. Nonetheless, it would be logical to think that the function of these proteins was annotated in cadmium-related experiments and that they could also be involved in a broader spectrum of stress response processes. In fact, there are studies confirming the overlapping response of plants to cadmium ion and other stress factors such as cold (Sergeant et al., 2014). Specifically, proteins like aldo-keto reductase 4 or formate dehydrogenase, which were linked to cadmium ion responses in the GO analysis, are known to take part in other stress events related to temperature (Turóczy et al., 2011; McNeilly et al., 2018).

After cross-validation of the results by comparing multivariate and univariate statistical analyses, 53 proteins were selected as top-significant. Some of them were involved in different phases of the life cycle of proteins, such as protein synthesis (ribosomal proteins and casein kinase II subunit alpha-1), folding (peptidyl-prolyl cis-trans isomerase CYP19-2), transport (mitochondrial import inner membrane translocase subunit TIM9) and degradation (proteases, proteasomes and deubiquitinating enzymes). These results are similar to the ones obtained at subsequent stages of the SE process when analysing the proteome of somatic embryos coming from EMs initiated under high temperatures (Castander-Olarieta et al., 2021a). Apart from the high fold changes observed for most ribosomal proteins, it is also highlightable the differential expression of several subunits of the 26S proteasome. This molecule, in addition to being a key element in the ubiquitin-mediated degradation pathway of proteins (Voges et al., 1999), has also been linked with several stress response processes in *A. thaliana* (Kurepa et al., 2008) and tobacco (Bahmani et al., 2017), and its interplay with cytokinin-mediated processes has also been documented (Smalle et al., 2002). This fact results of special interest as our previous proteomic study in somatic embryos also revealed a possible interconnection between the translasome machinery and cytokinins under high temperatures (Castander-Olarieta et al., 2021a). Moreover, other studies in EMs of radiata pine reported that the temperature during initiation of SE influences the profile of different types of cytokinins (Castander-Olarieta et al., 2021b, 2021c).

In opposition to our previous work in somatic embryos, few HSPs and chaperones were detected as top-significant in this study, and their expression was significantly reduced under high temperatures. On the contrary, a great variety of proteins directly or indirectly related to oxidative stress responses were selected. Even if most of them presented a down-regulation tendency, such as ubiquinol oxidase 1a, whose function as alternative oxidase is known to be important during both biotic and abiotic stress (Vanlerberghe, 2013), three proteins were up-regulated in response to heat: formate dehydrogenase, thioredoxin H-type and glutathione S-transferase DHAR3. As previously mentioned, formate dehydrogenase is a universal stress protein involved in the responses to various abiotic and biotic stresses by catalysing the oxidation of formate, a byproduct of ROS scavenging, and maintaining a reducing environment in the cell (Kurt-Gür et al., 2018). Several studies have demonstrated that heat can induce the synthesis of this enzyme (Ren et al., 2019) and its abundance seems to be regulated by the 26S proteasome (McNeilly et al., 2018). Thioredoxin H is part of the thioredoxin antioxidant system in plants and its accumulation under high temperatures has been reported in *Populus euphratica*, among others (Ferreira et al., 2006). Interestingly, this enzyme is able to regulate the synthesis of certain amino acids such as isoleucine (Lemaire et al., 2004). In this regard, ketol-acid reductoisomerase, one of the key enzymes for the synthesis of isoleucine, was found at higher concentrations in EMs induced under high temperatures in this study. In parallel, previous studies carried out in our laboratory have confirmed that heat at initiation stage of SE increases the levels of this amino acid in radiata pine EMs (Castander-Olarieta et al., 2019). These results reaffirm the hypothesis postulated in our previous work in which we suggested that HSPs and chaperones are important during long-term heat stress responses and thermomemory acquisition, while proteins and metabolites involved in oxidative stress defence are required during earlier response stages.

Following the same pattern described for somatic embryos, most of the enzymes from central metabolism, including the glycolytic and the pentose phosphate pathways and the tricarboxylic acid cycle, were down-regulated in EMs initiated under high temperatures, indicating a reduction of carbon flux in those pathways. Particularly, the enzyme enolase seems to be highly responsive to high temperatures and could be a potential molecular marker of heat stress response, as it presented high fold changes in both EMs and somatic embryos (Castander-Olarieta et al., 2021a). However, its role and behaviour during these processes remain unclear because in other studies with different tree species like poplar the genes encoding this protein were considerably up-regulated under high temperatures (Ren et al., 2019). Curiously, only one enzyme involved in the pentose phosphate pathway was found at higher levels under heat conditions: transketolase-1. Transketolases are responsible for the synthesis of sugar phosphate intermediates during the reductive and oxidative pentose phosphate pathways, but potentially due to their relationship with thiamine, they have been proposed as important enzymes during stress response (Rapala-Kozik et al., 2012).

In line with these findings, changes were also observed for the levels of proteins involved in the synthesis of cell wall cellulosic and non-cellulosic polysaccharides (cellulose synthase-like protein D3, UDP-arabinopyranose mutase 1), in the synthesis of starch and in the synthesis, elongation and transport of fatty acids (3-oxoacyl-[acyl-carrier-protein] synthase I, mitochondrial dicarboxylate/tricarboxylate transporter DTC). These results support the idea that thermotolerance acquisition requires cell membrane stabilization and the action of cell wall remodelling enzymes to fine-tune cellulose-hemicellulose networks by the biosynthesis of specific polymers or *in muro* modifications (Le Gall et al., 2015). Similar to what observed in this study, heat stress led to a general decrease in the sugar concentrations of both hemicellulose and cellulose fractions in corn plants (Suwa et al., 2010), whereas in coffee leaves, temperature had contrasting effects depending on the polymer fraction and specific sugars considered (Lima et al., 2013). Overall, it seems clear that heat can alter the structure and composition of cell walls, but its effect is both genotype and species-specific. Interestingly, alterations in sugar metabolism, such as changes in the carbon flux of the glycolytic and pentose phosphate pathways, and a reduction in the activity of enzymes involved in the synthesis of polysaccharides, were concomitant with increased levels of certain free soluble sugars like glucose and fructose under high temperatures, as confirmed by HPLC. This could be a side effect of those metabolic rearrangements, or an intentional drift towards the accumulation of small organic molecules that could act as compatible solutes under stress conditions (Ghosh et al., 2021). In this sense, Marias et al. (2017) and Matías et al. (2016) also detected higher levels of glucose and fructose in different conifer trees (*Abies alba, P. ponderosa, Pseudotsuga menziesii*) in response to elevated temperatures, presumably derived from starch hydrolysis.

Another interesting group of proteins differentially responding to heat treatments include structural constituents of the cytoskeleton (tubulin alpha-1 chain), chaperones required for the folding of cytoskeleton filaments (T-complex protein 1 subunit gamma) (Ahn et al., 2019) and proteins involved in cell patterning and cell division (Rop guanine nucleotide exchange factor 7, ribonucleoside-diphosphate reductase large subunit). All these proteins have interconnected functions, as well as a close relationship with cell-wall remodelling/organization proteins. Heat is known to disturb microtubular cytoskeleton assembly and organization, as observed in *A. thaliana* (Lei et al., 2020) and poplar (Wang et al., 2017), resulting in altered cytokinesis. On top of that, the effects of high temperatures on the cytoskeleton can thereby provoke alterations in the process of vesicle transport and cell wall deposition (Parrotta et al., 2016). Additionally, Rop guanine nucleotide exchange factor 7 is essential for a proper embryo patterning because of its involvement in the establishment and maintenance of the root quiescent centre (Chen et al., 2011). Recent research has demonstrated that this protein can also determine the structure of cell walls by directing the formation of cell wall pits through interaction with microtubules (Nagashima et al., 2018). Based on these results, we could hypothesise that the decrease in both initiation ad proliferation rates and the micromorphological changes observed in EMs subjected to heat stress in our previous studies (Castander-Olarieta et al., 2019) could be explained by the distinct profile of the aforementioned proteins.

Ultimately, we have to remark the differential accumulation of several proteins associated with epigenetic, transcription, and post-transcriptional regulation mechanisms under the different temperature conditions: s-adenosylmethionine synthase, zinc finger CCCH domain-containing protein 31, multiple organellar RNA editing factor 8 and splicing factor 3B subunit 6-like protein.

S-adenosylmethionine synthase catalyses the synthesis of s-adenosylmethionine from methionine and adenosine triphosphate, which is the central constituent of the methionine cycle and serves as methyl donor for DNA and histone methyltransferases, among other molecules. Consequently, alterations in the levels of this enzyme could provoke modifications in the DNA methylation status of cells, as observed in Li et al. (2011) and Lindermayr et al. (2020). Even if S-adenosylmethionine synthase is involved in a wide variety of methylation mechanisms, our results and the ones from previous works suggest that heat provokes long-lasting epigenetic changes in our SE model and S-adenosylmethionine synthase could take active part on them. In fact, radiata pine somatic embryos developed from EMs initiated at hight temperatures showed distinct levels of the enzyme responsible for the synthesis of adenosylhomocysteine, the competitive inhibitor of s-adenosylmethionine-dependent methyl transferase reactions (Castander-Olarieta et al., 2021a), and somatic plants produced under the same temperature conditions showed altered methylation status (Castander-Olarieta et al., 2020).

In the same way, zinc finger CCCH domain-containing protein 31 belongs to the CCCH zinc finger protein family, which can act as transcription factors regulating the expression of genes involved in plant development and stress response (Pi et al., 2018). Recent research has demonstrated that these proteins can also bind RNA and co-localize with processing bodies and stress granules, which play important roles in post-transcriptional regulation and epigenetic modulation of gene expression (Bogamuwa and Jang 2016).

Changing profiles of multiple organellar RNA editing factor 8 and splicing factor 3B subunit 6-like protein suggest alterations in RNA processing. Multiple organellar RNA editing factor 8, also known as MORF, is a pivotal factor of the RNA editosome by performing conversion editing events, mainly C-to-U. This protein seems to have a relevant role during stress response processes, as observed in poplar (Wang et al., 2019). However, the nature of RNA editing processes in plants is still unsure. At many sites, RNA editing is essential for expressing functional proteins and could be observed as a repair mechanism, whereas at other sites, RNA editing could serve as a regulatory process of gene expression (Chateigner-Boutin and Small 2010). Splicing factor 3B is involved in both constitutive and alternative splicing of precursor mRNAs. This last phenomenon is essential to enhance transcriptome plasticity and proteome diversity and is known to be a critical post-transcriptional regulation mechanism during diverse stress-response processes (AlShareef et al., 2017; Ling et al., 2017). Furthermore, the latest research carried out on this topic points towards a crosstalk among alternative RNA processing and epigenetic regulation upon stress response in plants (Zhang et al., 2020). These authors also suggest a possible feedback regulation mechanism of RNA processing on epigenetic mechanisms, including DNA methylation.

In summary, this work revealed new insights into thermopriming phenomenon in radiata pine EMs at the proteome and metabolic level. Similar to what observed at later stages of the SE process, high temperatures led to a deep reorganisation of the machinery required for protein synthesis, folding, transport and degradation, with special emphasis put on the 26S proteasome. Heat also provoked a general reduction of central metabolism enzymes and chaperones, whereas the levels of proteins with antioxidant activity and proteins involved in the synthesis of isoleucine were notably increased under high temperatures. This experimental setup enabled the identification of metabolic changes associated with cell patterning, cell division and synthesis of cytoskeleton polymers and cell wall polysaccharides, resulting in altered profiles of soluble sugars under the different temperature conditions. This work also reinforced our previous results in which we suggested that different epigenetic, transcriptional and post-transcriptional regulation mechanisms are underlying thermopriming and thermomemory acquisition and maintenance.

## Author contributions

PM, IM, JC, SC, CP, and AC-O conceived and planned the experiments. AC-O prepared all the plant material and executed the heat stress experiments. AC-O, CP, VM, and SC carried out the protein analysis. AC-O, CP, and SS-Á performed the soluble sugar content quantification. AC-O, VM, SC, and BM carried out the data curation and statistical analysis. AC-O wrote the manuscript. All authors provided critical feedback and helped to shape the research, analyses, and manuscript

