## Supplementary Figure 1 for "Thermopriming-associated proteome and sugar content responses in *Pinus radiata* embryogenic tissue"

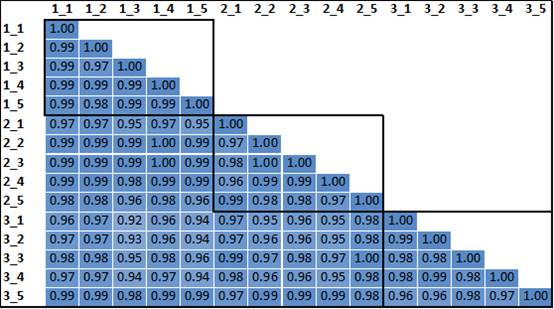

Supplementary Figure 1. Correlation analysis between samples using the Analysis ToolPak from Excel and the Pearson function. The first number of each pair indicates the temperature treatment, being 1 = Cond1, 2 = Cond2 and 3 = Cond3, whereas the second one refers to the sample number. The black boxes group the correlation values obtained among samples from the same temperature treatment.
